# A new lipid force field (FUJI)

**DOI:** 10.1101/373183

**Authors:** Nozomu Kamiya, Keiko Shinoda, Hideaki Fujitani

## Abstract

To explore inhomogeneous and anisotropic systems such as lipid bilayers, the Lennard-Jones particle mesh Ewald (LJ-PME) method was applied without a traditional isotropic dispersion correction. As the popular AMBER and CHARMM lipid force fields were developed using a cutoff scheme, their lipid bilayers unacceptably shrank when using LJ-PME method. A new lipid force field (FUJI) was developed on the basis of the AMBER force field scheme including the Lipid14 van der Waals parameters. Point charges were calculated by the restrained electrostatic potentials of many lipid conformers. The torsion energy profiles were calculated by high level *ab initio* molecular orbitals (LCCSD(T)/Aug-cc-pVTZ//LMP2/Aug-cc-pVTZ); then, the molecular mechanical dihedral parameters were derived by means of a fast Fourier transform. Incorporating these parameters into a new lipid force field without any fittings to experimental data, desirable lipid characteristics such as the area per lipid and lateral diffusion coefficients were obtained by GROMACS molecular dynamics simulations using the LJ-PME method and hydrogen virtual sites. The stability and structures of large membranes with undulatory fluctuations were studied by a multidrug efflux transporter (AcrABZ-TolC) with inner and outer membranes.

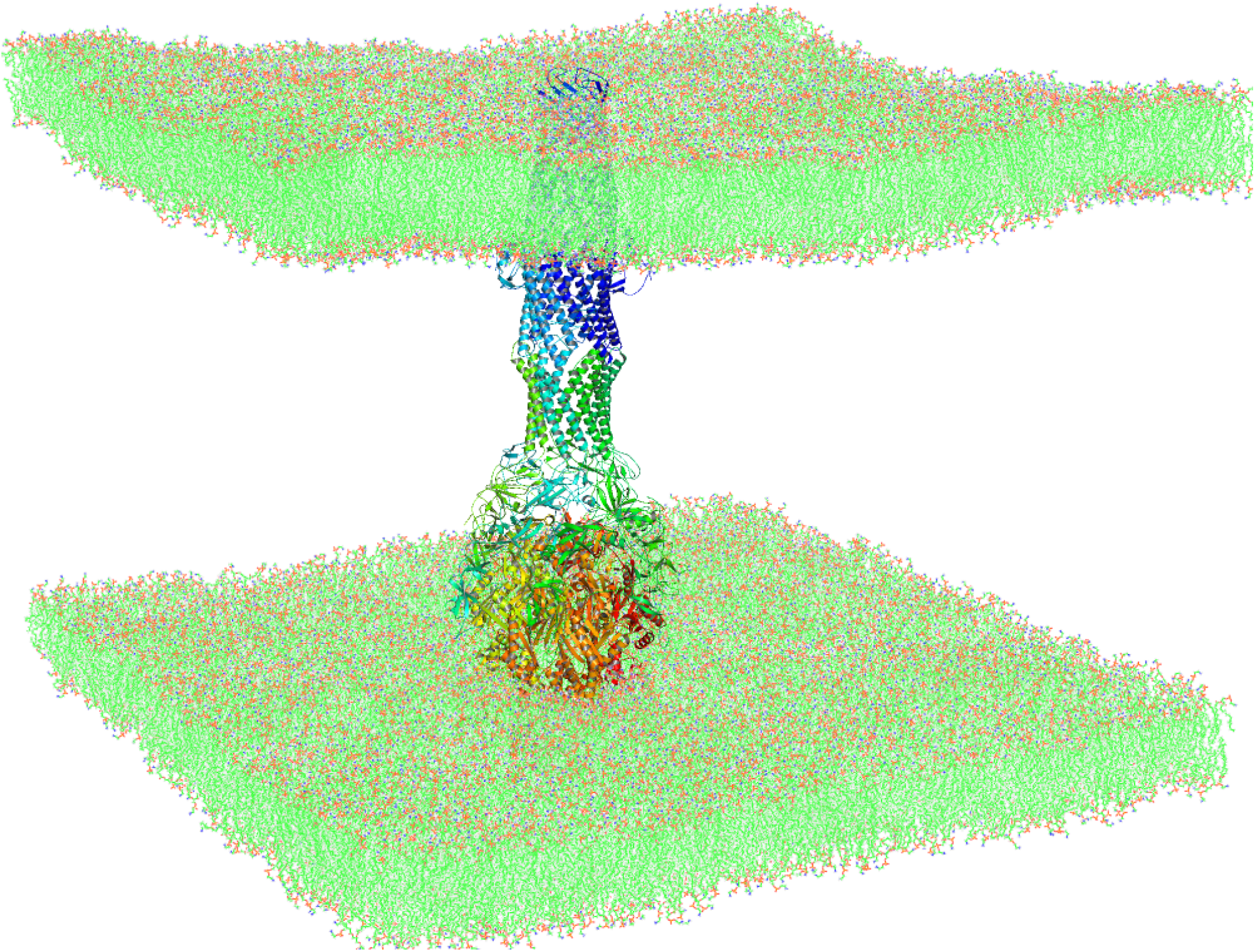

## 1 Introduction

Lipid bilayers separate the inside living part of a cell from the outside surroundings. Membrane proteins such as G-protein coupled receptors (GPCRs) can function properly with appropriate lipid bilayers.^1^ In order to investigate membrane proteins, we should have reasonable models of the lipid bilayers, which have two-dimensional structures with different characteristics in the perpendicular and longitudinal directions. The nonbonding interactions of molecules consist of Coulomb and van der Waals interactions within the scheme of classical molecular mechanics. Ewald summation was developed as a method for calculating the electrostatic energy of periodic systems such as ionic crystals.^2^ In molecular dynamics (MD) simulation for biomolecules, the particle mesh Ewald (PME) method was introduced to take into account the long-range characteristics of Coulomb interactions as an *N* ln(*N*) scale method.^3^ The van der Waals interaction consists of short-range core repulsion and quantum dispersion attraction. The repulsion has an exponential or *r*^−12^ functional form, which rapidly decays in comparison with the dispersion interaction of *r*^−6^. In comparison with the Coulomb interaction, the van der Waals interaction has a short-range characteristic; thus, a cutoff in the range of 10–14 Å was commonly used with the Buckingham or Lennard-Jones functions. However, the discarded dispersion attraction outside the cutoff still significantly affects the system, whose energy and pressure should be corrected to reproduce the proper values. The dispersion correction was formulated in an analytical and homogeneous manner with the averaged Lennard-Jones parameters of the van der Waals interaction.^4,5^

Using the electrostatic PME and Lennard-Jones cutoff schemes with the isotropic dispersion correction, lipid force fields such as GAFFlipid,^6^ Slipid,^7^ Lipid14,^8^ and CHARMM36 lipids^9^ were developed, which properly reproduced the experimental values of the surface area per lipid without additional surface tensions. However, this scheme applied the isotropic correction to basically inhomogeneous and anisotropic membrane systems. Wennberg et al. developed the Lennard-Jones PME (LJ-PME) method in order to improve the precision and accuracy of membrane simulations.^10^ They confirmed that the long-range contributions were well-approximated by the dispersion corrections in simple systems, but there were large effects on the surface tension for lipid bilayers, resulting in up to 5.5% deviation in the area per lipid and order parameters.

The AMBER and CHARMM force fields use the Lorentz–Berthelot (LB) combination rule to derive the Lennard-Jones distance parameter *σ_ij_* and potential parameter є*_ij_* as

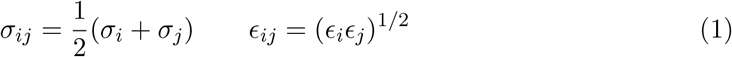

The LB combination rule requires the expensive calculations of several separate fast Fourier transforms in the implementation of the LJ-PME method. Using geometric combination rules in reciprocal space with modified interactions in direct space, Wennberg et al. developed an accurate and efficient high-performance LJ-PME method,^11^ which was implemented in the GROMACS MD package.^12,13^ Around room temperature, the chemical bonds in biomolecules can be constrained as a sound approximation of a quantum oscillator. Moreover, the angular vibrations involving hydrogen atoms can be removed using GROMACS virtual interaction sites. These constraints result in an additional advantage for MD simulations, which enables a longer time step up to 5 fs.^14^ When we take into account the more accurate effects of a system’s anisotropy and quantum oscillation by the LJ-PME and vibration constraints, the current lipid force fields give unsatisfactory results.

The FUJI force field was developed to describe the arbitrary organic molecules of proteins and small molecules in a unified manner,^15^ starting from the first version of the general AMBER force field (GAFF).^16^ We redefined the GAFF atom types and refined the force field parameters by a first-principles theoretical method, in contrast to the common empirical fitting to experimental data. For example, the torsion parameters of a protein backbone were determined to agree with the torsion energy profiles for model systems of a protein backbone calculated by high-level quantum mechanical theories such as the coupled-cluster method.^17^ The conformation preferences of dipeptides in water were measured by infrared and Raman spectroscopy,^18,19^ which revealed short-lived states in comparison with nuclear magnetic resonance (NMR) spectroscopy. Various force fields were compared with the measured conformation preferences, and the FUJI force field was the best.^20,21^

In this work, we developed a new lipid force field based on the same principles as the FUJI force field, and its characteristics were examined by MD simulations with the LJ-PME method and hydrogen virtual sites for pure lipid bilayers and large membrane systems with a multidrug efflux transporter (AcrABZ-TolC).

## 2 Lipid force field refinement

Phospholipids are amphiphilic with a hydrophilic (water-loving) phosphate head part and lipophilic (fat-loving or hydrophobic) fatty acid tails. They are long flexible molecules and show a variety of conformations in water. The lipid characteristics depend on the point charges, Lennard-Jones parameters, and torsion energy profiles of the lipid backbone. We introduced new atom types (Table 1) to retain the consistency of the FUJI force field and refined the force field parameters of phosphatidylcholine (PC) and phosphatidylethanolamine (PE), which have five types of fatty acid tails: lauroyl (LA), mylistrol (MY), oleoyl (OL), palmitoyl (PA), and stearoyl (ST).

**Table 1:** New atom types for the FUJI force field. The original GAFF atom types are on the right side in capitals.

|  |  |  |
| --- | --- | --- |
| Cox | carbonyl sp2 carbon in lipid | C |
| CTx | sp3 carbon in lipid tail | C3 |
| HAx | H bonded to sp2 carbon in lipid tail | HA |
| HCx | H bonded to aliphatic carbon in lipid tail | HC |
| N4x | sp3 N with four connected atoms in lipid | N4 |
| OPx | oxygen with one connected phosphorus in lipid | O |
| OSx | ether oxygen in lipid | OS |
| OTx | ether oxygen bonded to phosphorus in lipid | OS |
| Ox | oxygen with one connected carbon in lipid | O |

### 2.1 Point charges

Using MD simulations with the CHARMM36 lipid force field,^9^ we explored a variety of conformations of six PCs (DLPC: dilauroyl-PC, DMPC: dimyristoyl-PC, DOPC: dioleoyl-PC, DPPC: dipalmitoyl-PC, POPC: palmitoyl-oleoyl-PC, and SOPC: stearoyl-oleoyl-PC). For a lipid molecule in water, we saved the lipid coordinates at 1 ns intervals, obtained 400 conformations of every lipid, and then performed HF/6-31+G(d) calculations in vacuum for structure optimization with the Gaussian09 package.^22^ Figure 1 (a) shows the energy distribution of the POPC optimized conformations. The energies extend from 0 to 18 kcal/mol. We selected 256 lower energy conformations and performed restrained electrostatic potential (RESP) charge calculations^23^ by an R.E.D. Perl script (version III 52)^24^ with HF/6-31+G(d) for the six PCs. Figure 1 (b) shows the root mean square deviations (RMSDs) of the RESP charges of the low energy conformations compared with those of the 256 conformations for POPC. The RMSDs of the 68 and 128 conformations are 0.006 and 0.003, respectively. If we increased the conformation sampling from 256 to 512, we supposed that the RMSD would be less than 0.001. We concluded that the RESP charges were sufficiently converged by the 256 conformation sampling.

**Figure 1:**
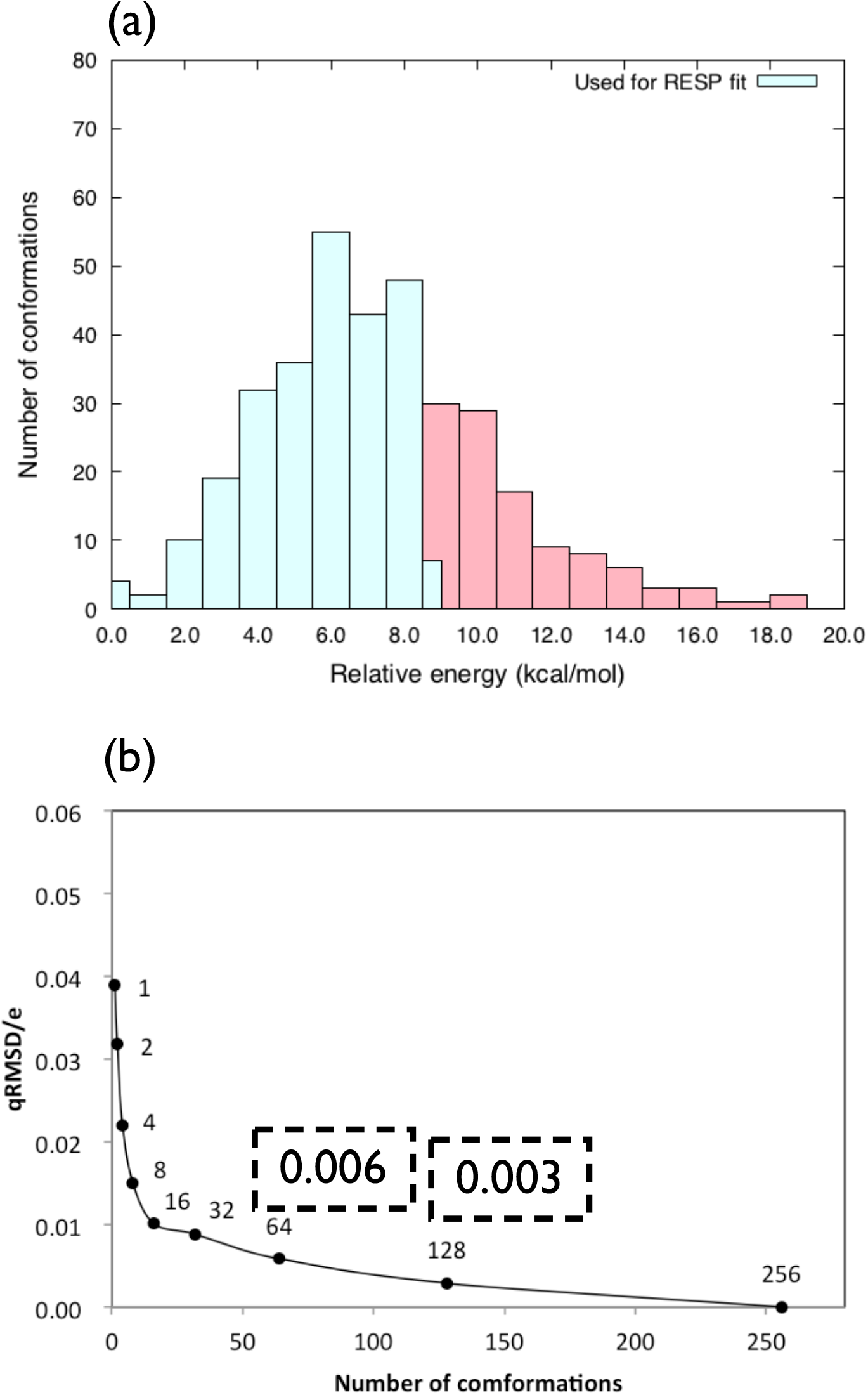
RESP charge calculations. (a) Energy distribution of the conformations after structural optimization by HF/6-31+G(d) calculations in vacuum. The light blue regions are the 256 conformations used for the RESP calculation. (b) Convergence of root mean square deviations (RMSDs) of the RESP charges.

The RESP charges near the head–tail connection part of DLPC are listed in Table 2. We use the CHARMM36 lipid atom names (Figure S1).^9^ The hydrocarbon charges obtained by R.E.D. were strange: carbon atoms were positively charged, and hydrogen atoms were negatively charged in the tail part. These are opposite signs to the common hydrocarbon, e.g., in the side chains of amino acids (alanine, leucine, etc.). To clarify the reason for this, we attempted different procedures. After obtaining the RESP charges for each conformation, we took the arithmetic averages of the 256 conformations, which are listed in the third column in Table 2. The charges of O21, C21, and O22 remained at the same level as those obtained by R.E.D. but the hydrocarbons showed smaller and noisy charges with positive and negative charges on the hydrogen atoms. We supposed that the noise originated from the amphiphilic characteristic of the lipid: large charges in the head part and tiny charges in the tail part. Therefore, we calculated the electrostatic potentials (ESPs) separately for the head and tail parts using the 256 conformations of the six PCs. The final point charges were obtained by the arithmetic averages of the RESP charges of all conformations of the head and tail parts. The final charges of O21, C21, and O22 exhibited very little change; moreover, the carbon atoms in the tail part were all negatively charged, and the hydrogen atoms were all positively charged, as indicated in the fourth column in Table 2. Compared with the Lipid14 charges, the tail hydrocarbon charges were much smaller, and C22 had less negative charge. We suppose that the C22 charge in Lipid14 includes some artificial errors caused by the RESP boundary condition between the head and tail parts.

**Table 2:** Point charges of DLPC near the head–tail connection part.

| Atom name | R.E.D. | Arithmetic<br>mean | Arithmetic<br>Head/tail | Lipid14 |
| --- | --- | --- | --- | --- |
| O21 | -0.4003 | -0.4038 | -0.4038 | -0.419523 |
| C21 | 0.7216 | 0.7611 | 0.7734 | 0.770445 |
| O22 | -0.5880 | -0.6074 | -0.6233 | -0.596883 |
| C22 | -0.0291 | -0.0486 | -0.0455 | -0.172159 |
| H2R | 0.0111 | 0.0180 | 0.0305 | 0.054418 |
| H2S | 0.0111 | 0.0180 | 0.0305 | 0.054418 |
| C23 | 0.0472 | 0.0202 | -0.0410 | 0.001510 |
| H3R | -0.0086 | 0.0055 | 0.0150 | 0.019108 |
| H3S | -0.0086 | 0.0055 | 0.0150 | 0.019108 |
| C24 | 0.0164 | -0.0081 | -0.0081 | -0.024078 |
| H4R | -0.0087 | 0.0060 | 0.0084 | 0.020580 |
| H4S | -0.0087 | 0.0060 | 0.0084 | 0.020580 |
| C25 | 0.0337 | 0.0002 | -0.0065 | -0.015735 |
| H5R | -0.0156 | 0.0003 | 0.0062 | 0.009551 |
| H5S | -0.0156 | 0.0003 | 0.0062 | 0.009551 |
| C26 | 0.0329 | -0.0001 | -0.0124 | -0.019631 |
| H6R | -0.0156 | 0.0004 | 0.0045 | 0.010236 |
| H6S | -0.0156 | 0.0004 | 0.0045 | 0.010236 |
| C27 | 0.0352 | 0.0074 | -0.0041 | -0.019279 |
| H7R | -0.0161 | -0.0018 | 0.0032 | 0.011207 |
| H7S | -0.0161 | -0.0018 | 0.0032 | 0.011207 |
| C28 | 0.0285 | 0.0014 | -0.0083 | -0.020879 |
| H8R | -0.0165 | -0.0023 | 0.0031 | 0.010385 |
| H8S | -0.0165 | -0.0023 | 0.0031 | 0.010385 |

Three methyl groups (CH_3_) of PC were replaced by the hydrogen atoms at the hydrogen equilibrium distance to the nitrogen atom (N); then, we obtained 256 conformations for each PE (DLPE, DMPE, DOPE, DPPE, POPE, and SOPE). From the ESP of the PE head part by HF/6-31+G(d), we calculated the RESP charges of each conformation to obtain the final RESP charges by the arithmetic average of all conformations. The final point charges of the head (PC and PE) and tail (LA, MY, OL, PA, and ST) parts are listed in the supplemental tables (Tables S1 and S2).

### 2.2 Lennard-Jones parameters

Many variants of the AMBER force field for proteins and nucleic acids (ff94,^25^ ff99,^26^ ff03,^27^ ff99SB,^28^ ff99SB-ILDN,^29^ and ff14SB^30^) and the first version of GAFF used the same Lennard-Jones parameter set. The FUJI force field uses GAFF atom types with capital letters for proteins and small molecules.^15^ Although newer versions of GAFF introduced modified Lennard-Jones parameters, the FUJI force field retained the use of the original AMBER Lennard-Jones parameters by optimizing the torsion dihedral parameters. However, for lipid modeling, it might be necessary to introduce new atom types with different Lennard-Jones parameters. Slipid introduced new lipid AMBER force field parameters, starting from the CHARMM36 lipid parameters.^7^ GAFFlipid introduced new Lennard-Jones and torsion dihedral parameters for the lipid tails.^6^ Lipid14 modified the Lennard-Jones and torsion dihedral parameters of both the head and tail groups.^8^ These lipid force fields successfully retained a reasonable density for the lipid bilayers without artificial surface tension control.

Table S3 compares the modified Lennard-Jones parameters of Lipid14 with the conventional AMBER parameters. For the lipid tail groups, Lipid14 modified the Lennard-Jones parameters of the hydrogen atoms but not those of the carbon atoms. Using methyl acetate to model the ester linkage, which connects the head and tail groups in a lipid, the oxygen and carbon parameters were modified. The modified oxygen parameters were also applied to the oxygen atoms in the phosphate group. With these minimum modifications of the Lennard-Jones parameters and some optimization of the torsion dihedral parameters, Lipid14 successfully produced excellent values for the area per lipid in bilayer simulations using a cutoff scheme for the van der Waals interaction with the isotropic dispersion correction. Since all of the Lipid14 modifications of the Lennard-Jones parameters weaken the van der Waals attractive interaction, they significantly contributed to the removal of the artificial surface tension term.

### 2.3 Torsion energy profiles

The accuracy of various levels of quantum mechanical theory was investigated to calculate the torsion energy profiles of the Ramachandran angles *ϕ* and *ψ* of the protein backbone in Ref. 17. Because of the accuracy and computational efficiency for calculating the energy profiles of the torsion angles in model systems, it was best to use the density-fitting (df) local correlation coupled-cluster singles and doubles with perturbative noniterative triples correction at the geometry optimized by the df local correlation second-order Møller– Plesset perturbation theory (MP2) with the Aug-cc-pVTZ basis set (df-LCCSD(T)/Aug-cc-pVTZ//df-LMP2/Aug-cc-pVTZ).^31^ In this work, we adopted the same method using the MOLPRO *ab initio* program package (version 2015.1).^32^

We used four model molecules to investigate the torsion energy profiles of PCs (Figure 2). Table 3 lists the torsion angles that were studied as representatives of the same atom type torsion angles. Since all of the torsion angles are symmetric, it was sufficient to calculate the energy profiles from 0*°* to 180*°*. Fixing the torsion angle at 15*°* intervals, the molecular structure was optimized to obtain the lowest total energy in vacuum. Since the optimized structure might depend on the initial structure, it was common to perform a couple of optimizations from different initial structures and determine the lowest energy structure at a fixed torsion angle. The dots in Figure 3 show the total energy profiles *E_QM_* (*ϕ*) around the torsion angle *ϕ* calculated by MOLPRO with df-LCCSD(T)/Aug-cc-pVTZ//df-LMP2/Aug-cc-pVTZ. Each dot has an *ab initio* optimized molecular structure.

**Figure 2:**
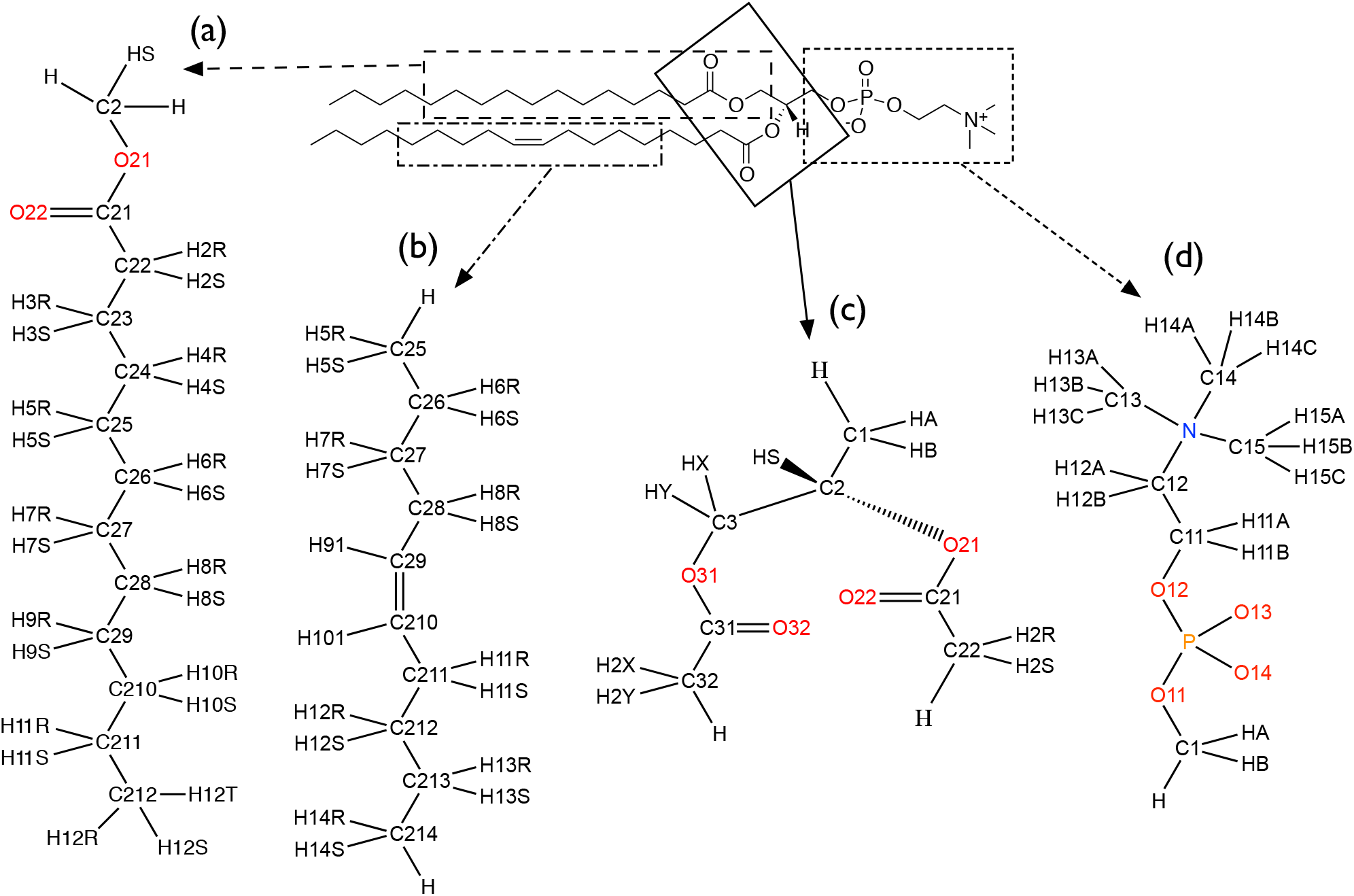
Four model molecules extracted from phosphatidylcholine to derive the torsion dihedral parameters (*V_n_*, *γ_n_*).

**Table 3:** Torsion angles in the four model molecules with atom names (Figure 2) and atom types.

| Model molecule | Atom name | Atom type |
| --- | --- | --- |
| (a) | C25-C26-C27-C28 | CTx-CTx-CTx-CTx |
|  | C21-C22-C23-C24 | Cox-CTx-CTx-CTx |
|  | O21-C21-C22-C23 | OSx-Cox-CTx-CTx |
|  | C2-O21-C21-C22 | C3-OSx-Cox-CTx |
| (b) | C28-C29-C210-C211 | CTx-C2-C2-CTx |
|  | C27-C28-C29-C210 | CTx-CTx-C2-C2 |
|  | C26-C27-C28-C29 | CTx-CTx-CTx-C2 |
| (c) | O21-C2-C3-O31 | OSx-C3-C3-OSx |
|  | C21-O21-C2-C3 | Cox-OSx-C3-C3 |
| (d) | O11-P-O12-C11 | OTx-P5-OTx-C3 |
|  | P-O12-C11-C12 | P5-OTx-C3-C3 |
|  | O12-C11-C12-N | OTx-C3-C3-N4x |

**Figure 3:**
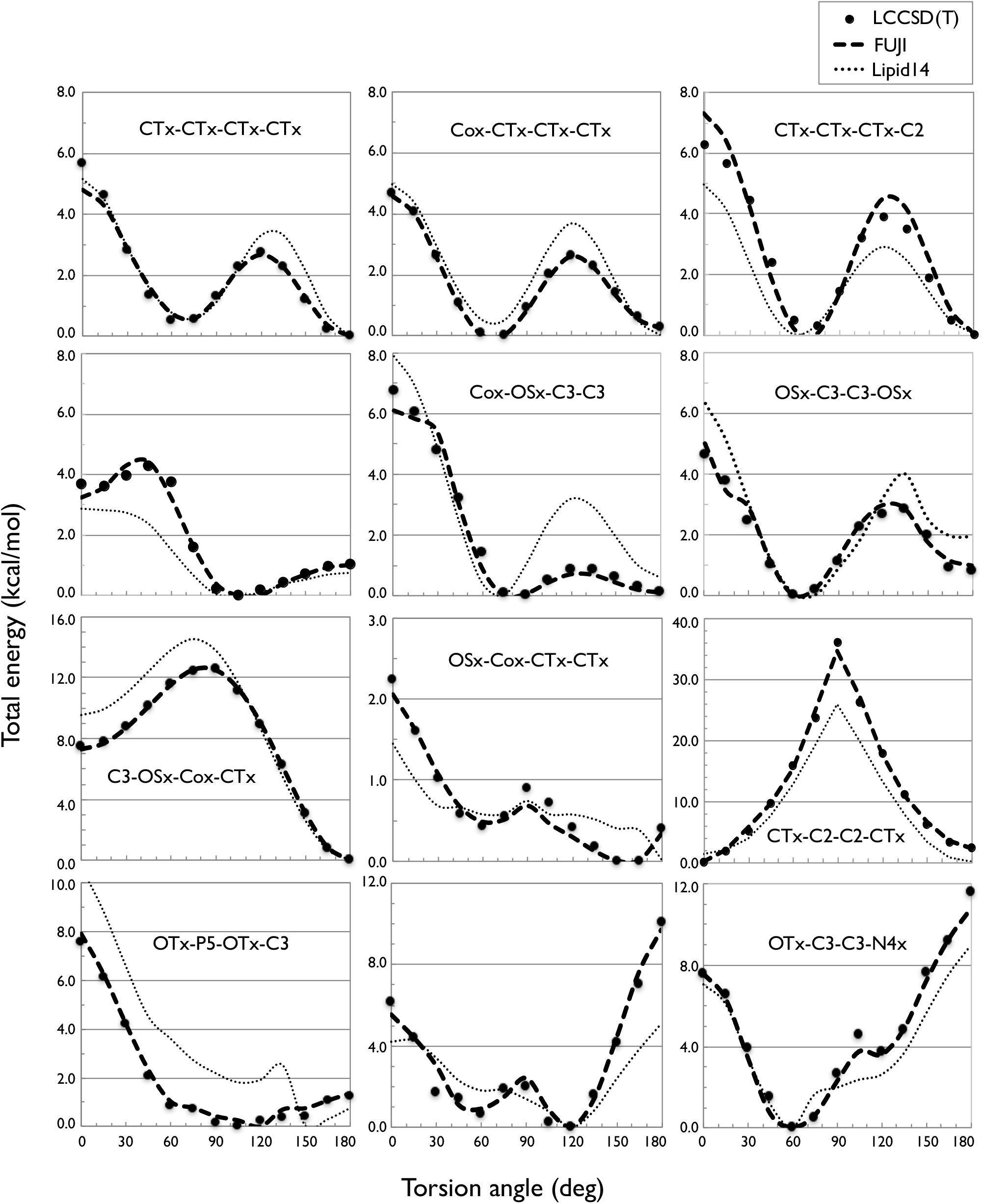
Torsion energy profiles of the 12 torsion angles in the four model molecules. The dots are the *ab initio* energies calculated by LCCSD(T)/Aug-cc-pVTZ//LMP2/Aug-cc-pVTZ. The thick dashed lines are the energy profiles obtained by the new lipid force field (FUJI). The thin dotted lines are the energies obtained by Lipid14 torsion dihedral parameters.

Figure 4 shows an example of a hurdle in obtaining the lowest energy structure at fixed torsion angles. The OTx-P5-OTx-C3 torsion angle in the model molecule (d) was fixed at 120*°* in the left and at 135*°* in the right. The terminal O12-P-O11-C1 torsion angle (Figure 2) was at 75*°* on the left and at −72*°* on the right. The terminal CH_3_ flipped owing to the 15*°* difference in the OTx-P5-OTx-C3 torsion angle. When we optimized the structure from the left structure fixing the OTx-P5-OTx-C3 torsion angle at 135*°*, the terminal O12-P-O11-C1 torsion angle became 76*°* and the total energy became 0.72 kcal/mol higher than the right structure. We needed other optimizations from different initial structures to obtain the lowest energy.

**Figure 4:**
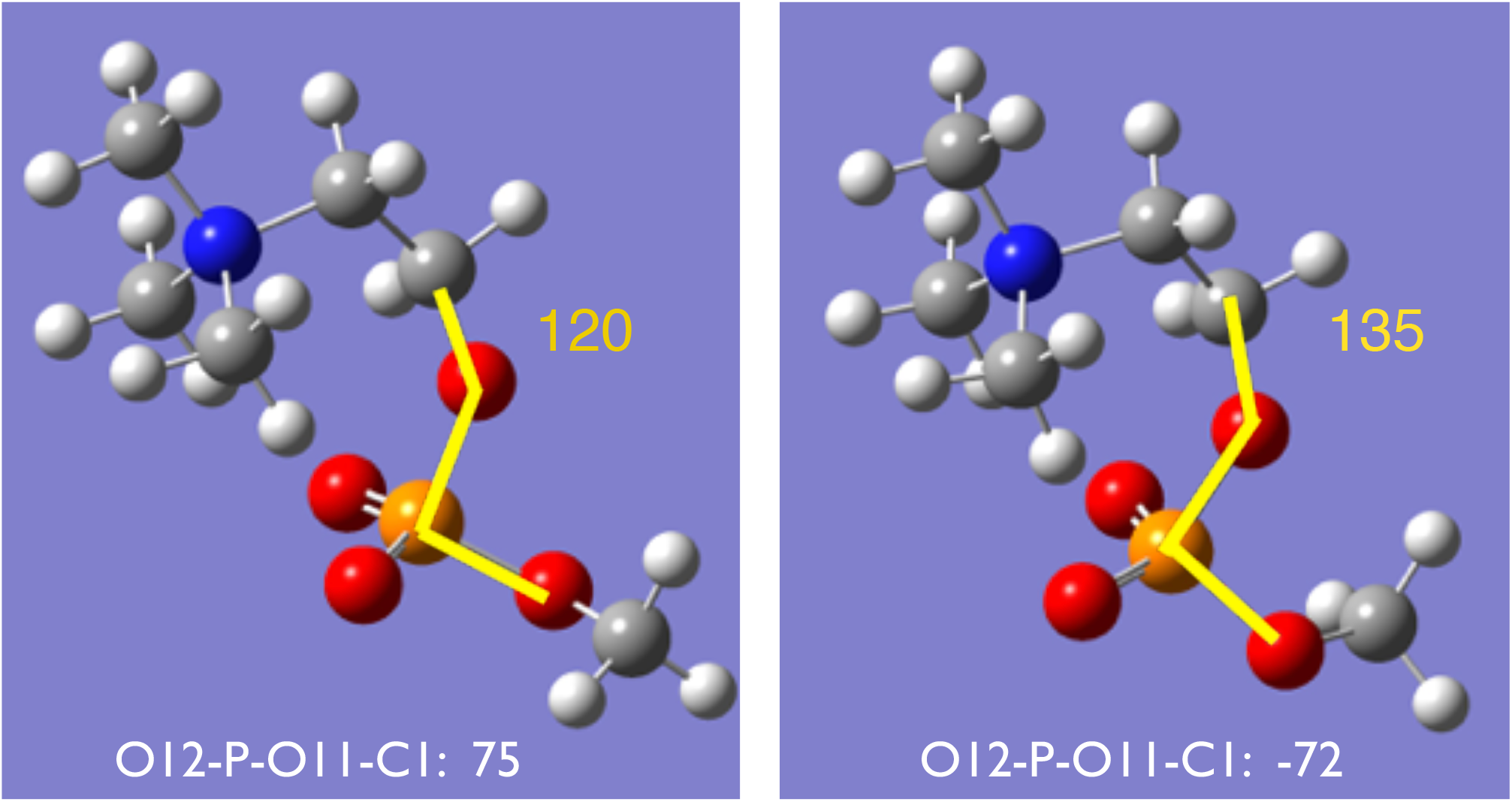
Optimized structures of model molecule (d): the OTx-P5-OTx-C3 torsion angle (yellow line) is at 120*°* on the left and at 135*°* on the right. The terminal CH_3_ is flipped by the O12-P-O11-C1 torsion angle.

The molecular mechanical total energy *E_MM_* for a conformation is given by

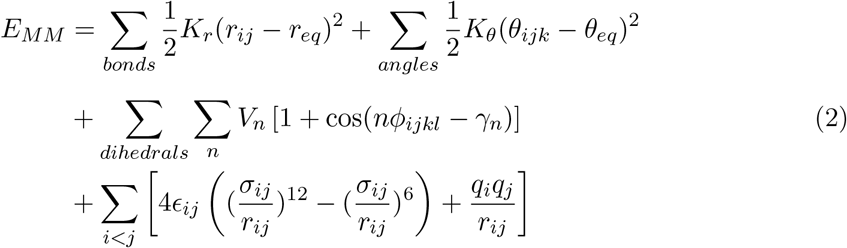

where *r_eq_* and *θ_eq_* are the equilibrium structural parameters and *K_r_* and *K_θ_* are force constants. *n* is the multiplicity, *γ_n_* is a phase angle, and *V_n_* is a force constant for the torsion energy. *є_ij_* and *σ_ij_* are the Lennard-Jones parameters, and *q_i_* is a point charge. We used the new RESP point charge for *q_i_* and the FUJI force field parameters for *r_eq_*, *θ_eq_*, *K_r_*, and *K_θ_*, which were used for proteins and small molecules depending on the atom types.^15^ We utilized two sets of Lennard-Jones parameters to derive new torsion dihedral parameters (*V_n_* and *γ_n_*): all conventional AMBER parameters or including the Lipid14 modifications.

We used GROMACS version 4.6.7 to derive new torsion dihedral parameters. For a target torsion angle *ϕ*, we derived 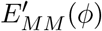 at each 15*°* interval, where the relevant force constants *V_n_* were set to zero and old force constants were used for the other torsion angles. Starting from the *ab initio* optimized structure, we performed molecular mechanical structure optimization with fixed *ϕ* at each 15*°* interval and then obtained 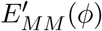. We defined *T_MM_* (*ϕ*) as the difference between the quantum *ab initio* energy *E_QM_* (*ϕ*) and the molecular mechanical energy 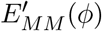:

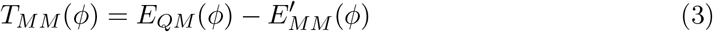

Since 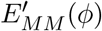 did not contain the relevant torsion energy, *T_MM_* (*ϕ*) should primarily consist of the relevant torsion energy. Since *E_QM_* (*φ*) and 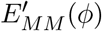 were defined from 0*°* to 180*°*, *T_MM_* (*ϕ*) could be expanded by a cosine Fourier series. We assumed that the finite cosine Fourier terms of *T_MM_* (*ϕ*) were the relevant molecular mechanical torsion energy. This means that *γ_n_* was restricted to 0*°* or 180*°*. Since *T_MM_* (*ϕ*) was defined at each 15*°* interval, the Fourier coefficients of *T_MM_* (*ϕ*) could be calculated by a fast Fourier transform.

The four model molecules had multiple target torsion angles (Table 3) whose torsion dihedral parameters should be derived to agree with the quantum *ab initio* energy *E_QM_* (*ϕ*). We conducted self-consistent derivations of multiple torsion dihedral parameters in a model molecule. First, using the model (a) molecule, we derived the new torsion dihedral parameters of C25-C26-C27-C28 by a fast Fourier transform to agree with the *ab initio* energies of CTx-CTx-CTx-CTx in Figure 3. The derived torsion dihedral parameters were applied to the same atom type torsion angles. Then, we derived the new torsion dihedral parameters of C2-O21-C21-C22, O21-C21-C22-C23, and C21-C22-C23-C24 sequentially to agree with the *ab initio* energies. We repeated this procedure a couple of times for the model molecule to obtain the final torsion dihedral parameters. The derived torsion dihedral parameters of CTx-CTx-CTx-CTx were applied to the model (b) molecule, and those of C3-OSx-Cox-CTx were applied to the model (c) molecule. In the same way, new torsion dihedral parameters were derived in the model (b), (c), and (d) molecules.

The thick dashed lines in Figure 3 show the molecular mechanical energies calculated by new torsion dihedral parameters and Lennard-Jones parameters with the Lipid14 modifications. Even if we did not include the Lipid14 Lennard-Jones modifications, we obtained almost the same dashed lines but slightly different torsion dihedral parameters by the same procedure. The thin dotted lines show the energies obtained with the Lipid14 torsion dihedral parameters. Table S4 lists our final torsion dihedral parameters, which were derived with the Lennard-Jones parameters including the Lipid14 modifications.

## 3 Lipid bilayers

Using the new force field parameters, we examined the bilayer characteristics of seven lipids (DLPC, DMPC, DOPC, DPPC, POPC, SOPC, and POPE). First, the lipid bilayer structures were constructed by the CHARMM-GUI membrane builder.^33^ Lipid topology files were generated by the GROMACS pdb2gmx subprogram with the hydrogen virtual site option. After system preparation (solvation, energy minimization, and thermal equilibration), we performed 300 ns MD simulations with the electrostatic PME and LJ-PME methods with geometric approximations of the combination rules in reciprocal space with 5 fs time steps and a linear constraint solver (LINCS) order of 6^14^ using the Verlet cutoff scheme. The simulation temperatures were controlled by the Nosè–Hoover method^34,35^ at different temperatures (293, 303, 323, and 333 K for DLPC, POPC and SOPC; 303, 323, and 333 K for DMPC; 288, 303, and 318 K for DOPC; 323, 333, and 353 K for DPPC; and 293, 303, 310, 323, and 333 K for POPE). During the first 200 ns, we used Berendsen pressure control^36^ and then used Parrinello–Rahman pressure control^37^ at 1 atm during the latter 100 ns. The unit cell of the bilayer contained 128 lipid molecules and 6400 TIP3P water molecules. We used GROMACS version 2016.3 with single precision for the bilayer simulations. We ran three simulations at each temperature and used the last 50 ns of the trajectories for analysis.

### 3.1 Area per lipid

Figure 5 shows the temperature dependence of the area per lipid for the seven lipids calculated from the dimensions of the simulation box in comparison with experimental values.^38–46^ The calculated values include the Lipid14 modifications of the Lennard-Jones parameters, and the standard errors in the three trajectories were between 0.1 and 0.4 Å^2^. When we utilized the original AMBER Lennard-Jones parameters, POPE gave an area per lipid of 47.8 Å^2^, in contrast to 58.6 Å^2^ at 303 K in Figure 5. The small increase in the attractive van der Waals interaction drastically shrank the bilayer, even though the torsion dihedral parameters were optimized for the original AMBER Lennard-Jones parameters. Although some experimental values were scattered, the results reasonably agreed with the experimental area per lipid. We concluded that the additional atom types with different Lennard-Jones parameters from the original AMBER ones in Table S3 were necessary to properly describe the lipid characteristics.

**Figure 5:**
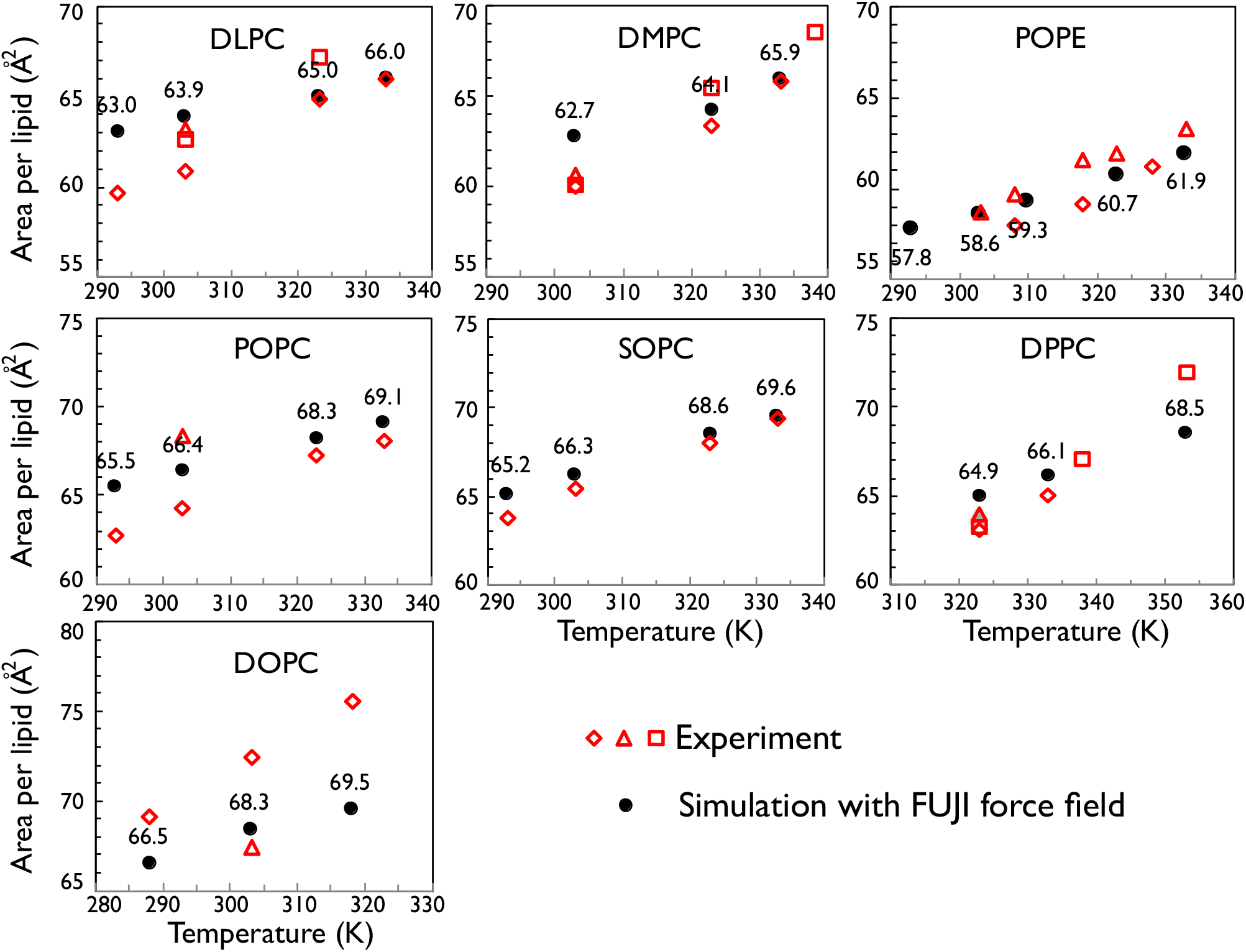
Area per lipid for the seven lipid bilayers. The black circles indicate simulation results. The red diamonds, triangles, and rectangles are experimental values (DLPC and DMPC from Refs. 38,41,45; DOPC from Refs. 43,44; DPPC from Refs. 38,39,45; POPC from Refs. 42,45; SOPC from Ref. 45; and POPE from Refs. 40,46).

### 3.2 Diffusion coefficient

Figure 6 shows the temperature dependence of the lateral diffusion coefficient of the seven lipid bilayers in comparison with experimental values.^47–53^ The simulation values were calculated by the GROMACS msd subprogram with a fitting time ranging from 10 to 20 ns and the option to remove the center of mass motion of the lipid bilayer. The experimental values were measured by different methods such as fluorescence recovery after photobleaching (FRAP),^47–49^ pulsed field gradient NMR (pfg-NMR),^51–53^ and resonance energy transfer (RET).^50^ The simulation results reasonably agreed with the experimental results. Our diffusion coefficients were slightly larger than those of Lipid14 (Table 9 in Ref. 8). Although the Lennard-Jones parameters are the same as those of Lipid14, the point charges differ. As our tail charges are smaller than the Lipid14 ones, this might reduce the attractive interactions between lipids so that the diffusion coefficients are larger.

**Figure 6:**
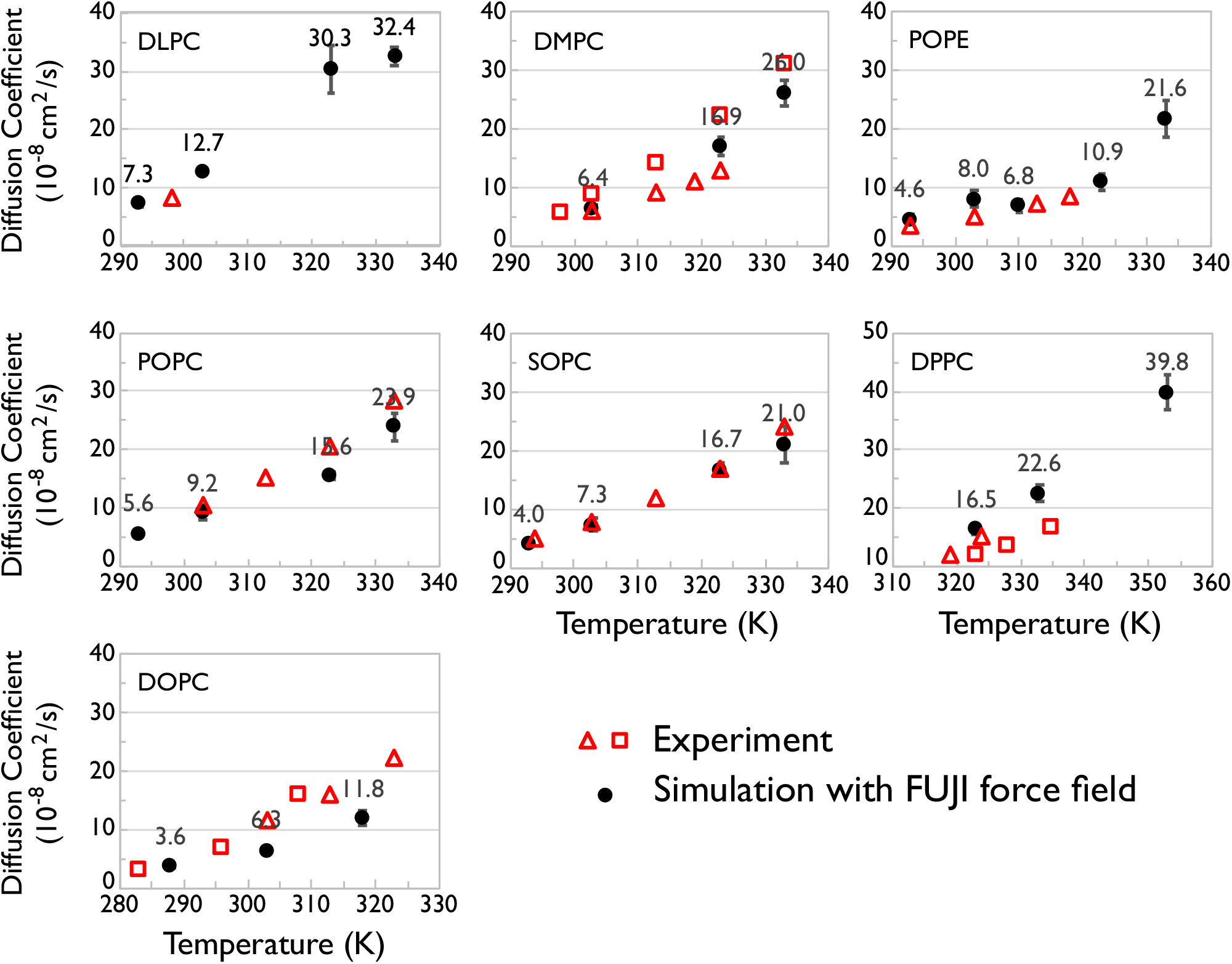
Lateral diffusion coefficients of the seven lipid bilayers. The black circles indicate simulation results, and the error bars indicate the standard errors in the three trajectories. The red triangles and rectangles are experimental values (DLPC from Ref. 49; DMPC from Refs. 48,51; DOPC from Refs. 50,51; DPPC from Refs. 47,52; POPC from Ref. 51; SOPC from Ref. 53; and POPE from Ref. 47).

### 3.3 Order parameter of the acyl tails

The order parameter of acyl chains is defined by

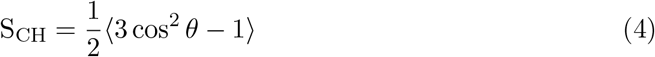

where *θ* is the angle between the bilayer normal (*z* axis in this case) and the C–H bond vector and 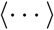 means the ensemble average. If all acyl tails are parallel to the bilayer normal with all trans C–C–C–C dihedrals, S_CH_ = −0.5, as *θ* = 0. This is a perfect ordered state. When the acyl tails were in disorder, *−*S_CH_ decreases from 0.5. Figure 7 shows our lowest temperature *−*S_CH_ versus the carbon position with the experimental *|*S_CH_*|* values. 38,54–56 When the temperature was increased, the *−*S_CH_ decreased so that the acyl tails became more disordered. Our calculated values of *−*S_CH_ were smaller than the experimental *|*S_CH_*|* values except those of DOPC, and also smaller than other simulation results by Lipid14 models,^8^ CHARMM36 lipid models,^9^ or Berger united-atom models.^55,57^ Although the united-atom model caused some discrepancy by the analysis tools for the acyl chain order parameter, many tools works well on all-atom models.^58^ We concluded that our results had more disordered acyl tails than the other models.

**Figure 7:**
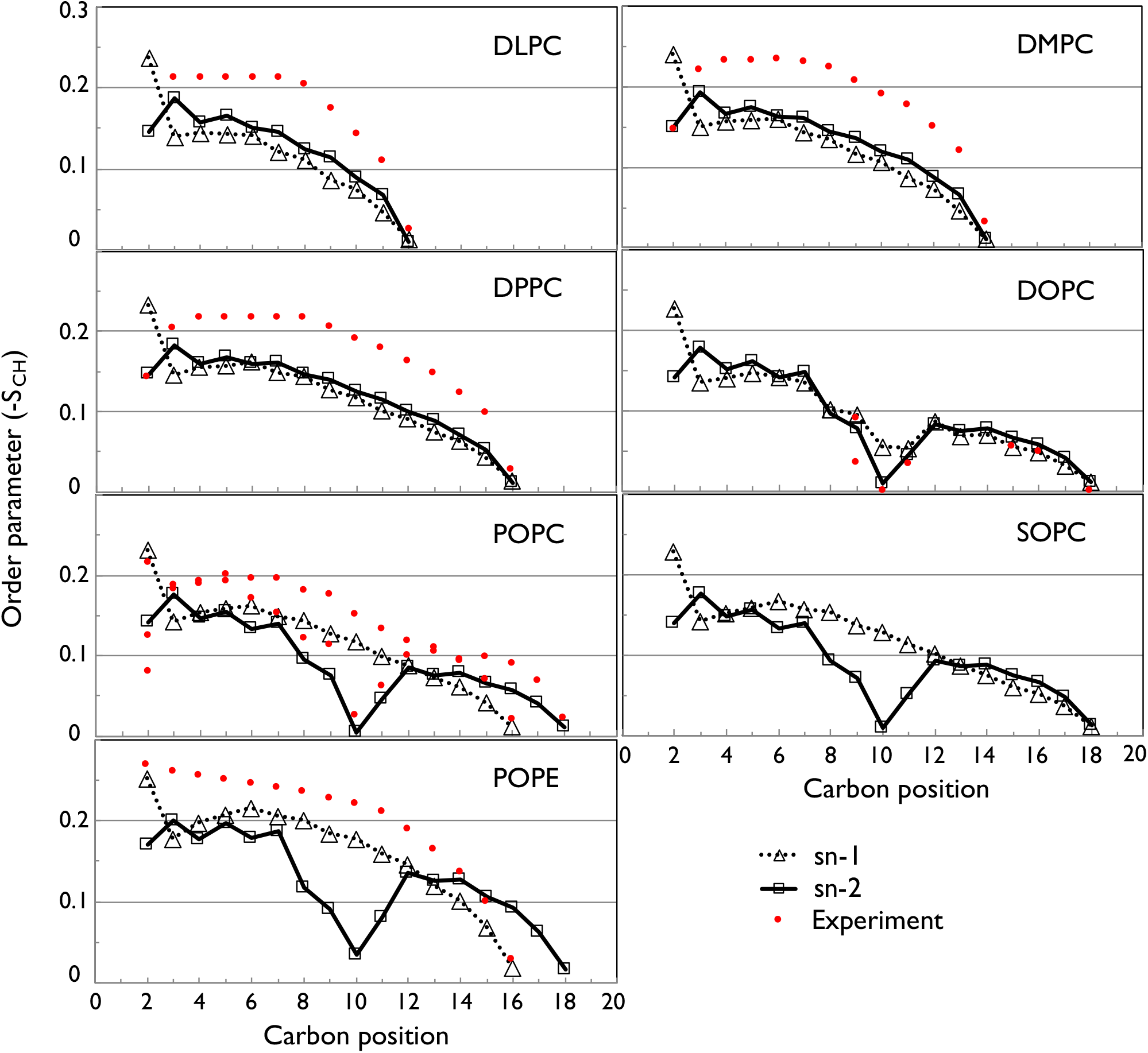
Calculated order parameter (–S_CH_) profiles for the acyl tails of the seven lipid bilayers with the NMR order parameter |S_CH_| taken from Ref. 38 for DLPC, DMPC, and DPPC; from Ref. 54 for DOPC; from Ref. 55 for POPC; and from Ref. 56 for POPE.

The order parameter of acyl chains is measured by NMR spectroscopy. Two types of NMR measure different physical observables: ^2^H NMR measures the quadrupolar splitting ∆*ν_Q_*, and H-^13^C NMR measures the effective dipolar coupling *d*_CH_. These NMR observables are related to the order parameter by different scaling factors. Although the scaling factors were determined by other experiments, some ambiguities remain.^59^ We scaled the 2H NMR values by 0.7 and the H-^13^C NMR values by 0.75 and plotted them with our results (Figure S2). We find satisfactory agreement.

### 3.4 Electron density and bilayer thickness

The electron density profiles along the lipid bilayer normal were calculated by the GRO-MACS density subprogram with the “-center” option.^60^ Figure 8 shows the electron density profiles derived from a trajectory at the lowest simulation temperature for each lipid bi-layer. Our results were similar to those of the Lipid14 model (Figure 8 in Ref. 8), although the order parameters of the acyl chains were different. From the electron density profiles, we derived the lipid bilayer thicknesses (D_HH_) from the peak-to-peak distance of the total electron density and compared with the experimental values. ^39–42,44,61^ They were in reasonable agreement.

**Figure 8:**
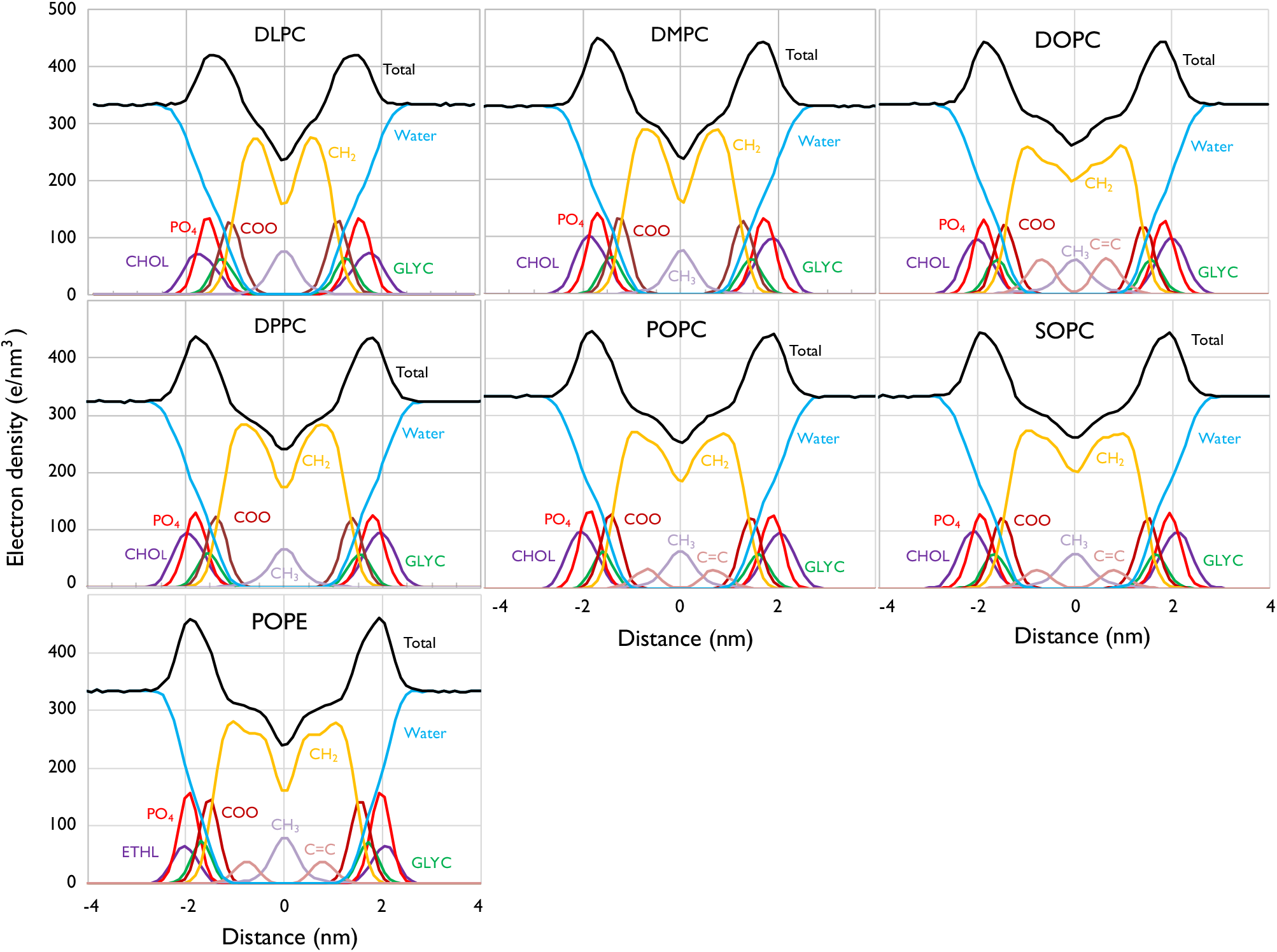
Electron density profiles along the membrane normals of the seven lipid bilayers. Each profile was calculated from a trajectory at the lowest simulation temperature. The profiles were decomposed into the groups choline (CHOL), phosphate (PO_4_), glycerol (GLYC), carbonyl (COO), methylene (CH_2_), unsaturated hydrocarbon (C=C), terminal methyl (CH_3_), and ethanolamine (ETHL).

## 4 Lipid bilayers with an AcrABZ-TolC efflux pump

Using small unit cells with 128 lipids, we obtained the structural parameters and compared them with the experimental values. However, it is well-known that large lipid bilayers exhibit fluctuations, where there are “waves” forming in the system. In order to compare with experiments, we might need to include the undulation smearing effect. To clarify the issue, we performed large bilayer simulations with membrane proteins. The multidrug efflux pump protein complex (AcrABZ-TolC) of *Escherichia coli* comprises three outer membrane proteins (TolC), six periplasmic membrane fusion proteins (AcrA), three inner membrane transporters (AcrB), and three small peptides (AcrZ), which modify the activity of AcrB. Using its cryo-electron microscopy (cryo-EM) structure (PDB code: 5NG5),^62^ we constructed an all-atom model of the whole AcrABZ-TolC pump with POPE inner and outer membranes. By MD simulations, we investigated the stability and atomic structures of the large lipid bilayers.

### 4.1 System setup and MD simulations

LAMBADA aligns the membrane and protein coordinates in the GROMACS GRO format by determining the protein’s hydrophobic belt directly exposed to the hydrophobic part of the lipid bilayer.^63^ Using the cryo-EM structure, we identified the inner and outer membrane positions by LAMBADA and removed the overlapping POPE molecules whose heavy atoms were within 0.12 nm of the heavy atoms of the protein. We added TIP3P water, Mg^2+^, Na^+^, and Cl– under charge neutrality conditions by the GROMACS subprograms solvate and genion.^64,65^ The system had a TolC trimer, an AcrA hexamer, AcrB and AcrZ trimers, 12,210 POPE molecules, 1,812,031 TIP3P molecules, 6,329 Na^+^ ions, 6,842 Cl^-^ ions, and 294 Mg^2+^ ions, and the total number of atoms was more than 7 million. The topology files of proteins and lipids were generated by the GROMACS pdb2gmx subprogram with the hydrogen virtual site option using GROMACS version 2016.4 and the FUJI force field files.

Using the electrostatic PME and LJ-PME methods, we minimized the energy of the system under the default criteria: “Fmax < 100” and a 2 ns MD simulation with the position restraint only on the heavy atoms of the protein and not on the heavy atoms of POPE at 298 K and 1 atm by the Nosè–Hoover thermostat and Berendsen barostat. Generating the initial velocities, we performed two 10 ns simulations without the restraint. Then, we performed two 400 ns MD simulations at 298 K by the Nosè–Hoover thermostat and at 1 atm by Parrinello–Rahman barostat with 5 fs time steps and an LINCS order of 6 using the Verlet cutoff scheme. During these procedures, there were no undesirable lipid behaviors such as the instability of the membrane structure, the escape of lipid molecules from the membrane, and so on.

Figure 9 shows short-range interaction energies of two trajectories (a) and (b). The short-range Lennard-Jones interactions (LJ-SR) were almost the same for the two trajectories, but the short-range Coulomb interactions (Coul-SR) slightly differed. The Coul-SR difference between TolC and AcrB is larger in trajectory (a) than in trajectory (b). In both trajectories, the Coul-SR between the membrane and TolC initially became stronger until 50 ns and then fluctuated in slightly different ways. Undulatory fluctuations of membranes were always observed in both trajectories. A snapshot of the structure at 300 ns for trajectory (a) is shown in Figure S3.

**Figure 9.**
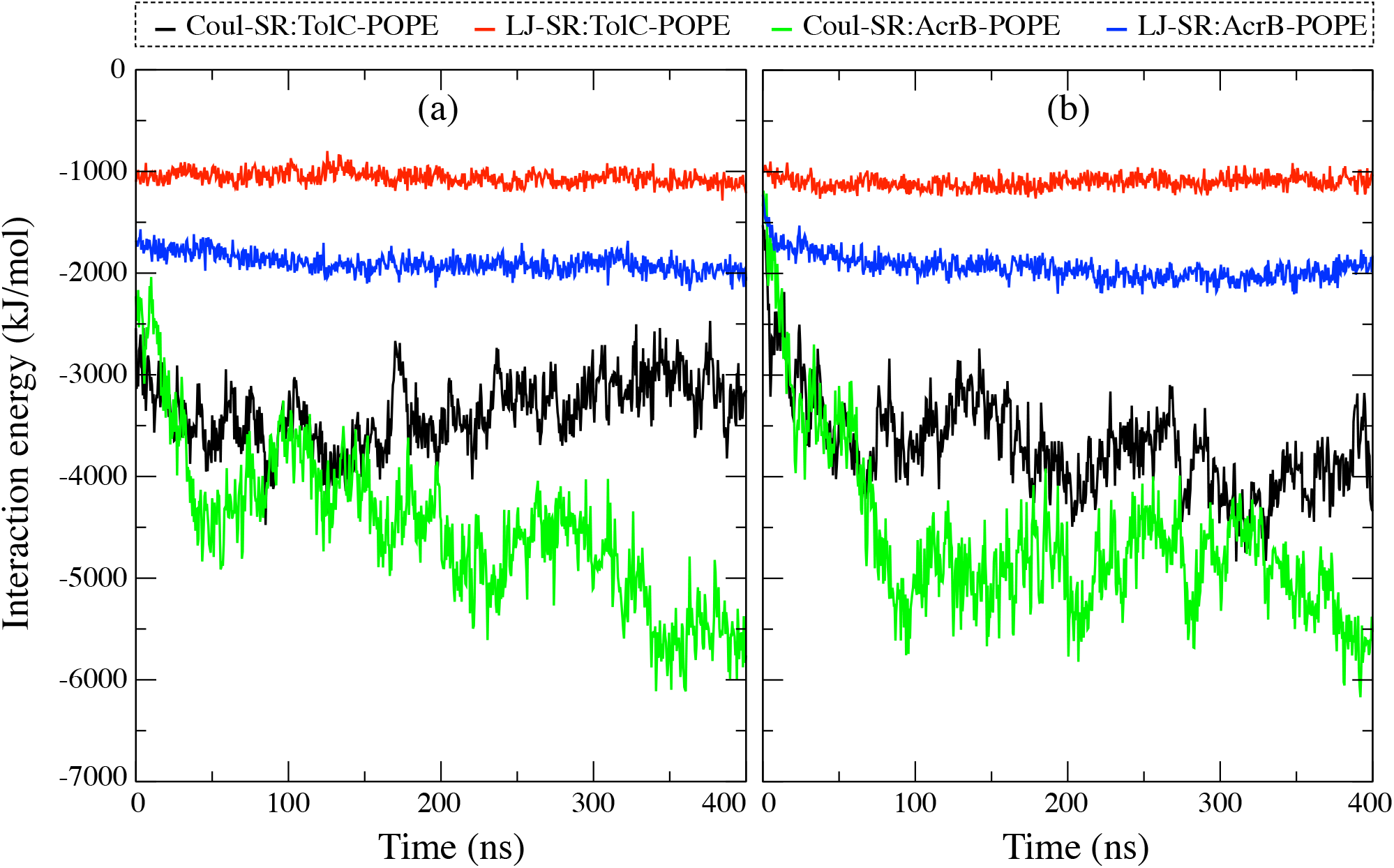
Short-range interaction energies of POPE with TolC and AcrB in the two 400 ns MD simulations.

### 4.2 Large membrane characteristics

The unit cell size of the pure POPE bilayer was a square area with a side length of about 6 nm times a height of 9 ns, and that of the AcrABZ-TolC membrane efflux system was a square area with a side length of 42 nm times a height of 39 nm. The size difference manifested the membrane undulatory fluctuations, whose shape changed roughly in a 10 ns time scale. According to the interaction energies in Figure 9, the initial structure relaxed until 50 ns. Using the trajectories from 50 to 400 ns, we calculated the electron density profiles of POPE and water in Figure 10 along the membrane normal by the GROMACS density subprogram. Since the proteins occupied some space, the absolute values of the electron density differed from the pure POPE bilayer (Figure 8). The profiles of the POPE parts were broader in the efflux system than in the small pure POPE bilayer. For example, the CH_2_ profile was within a thickness of 4 nm in the pure POPE bilayer, but the CH_2_ profiles in the efflux system spread in the range of more than 6 nm. We examined the order parameter of the acyl tails (*−*S_CH_) in the efflux system (Figure S4). Owing to the undulatory fluctuations, the POPE distributions significantly changed; however, the order parameters of the two trajectories were the same as that of the pure POPE bilayer, except for the effect due to a 5*°* temperature difference.

**Figure 10.**
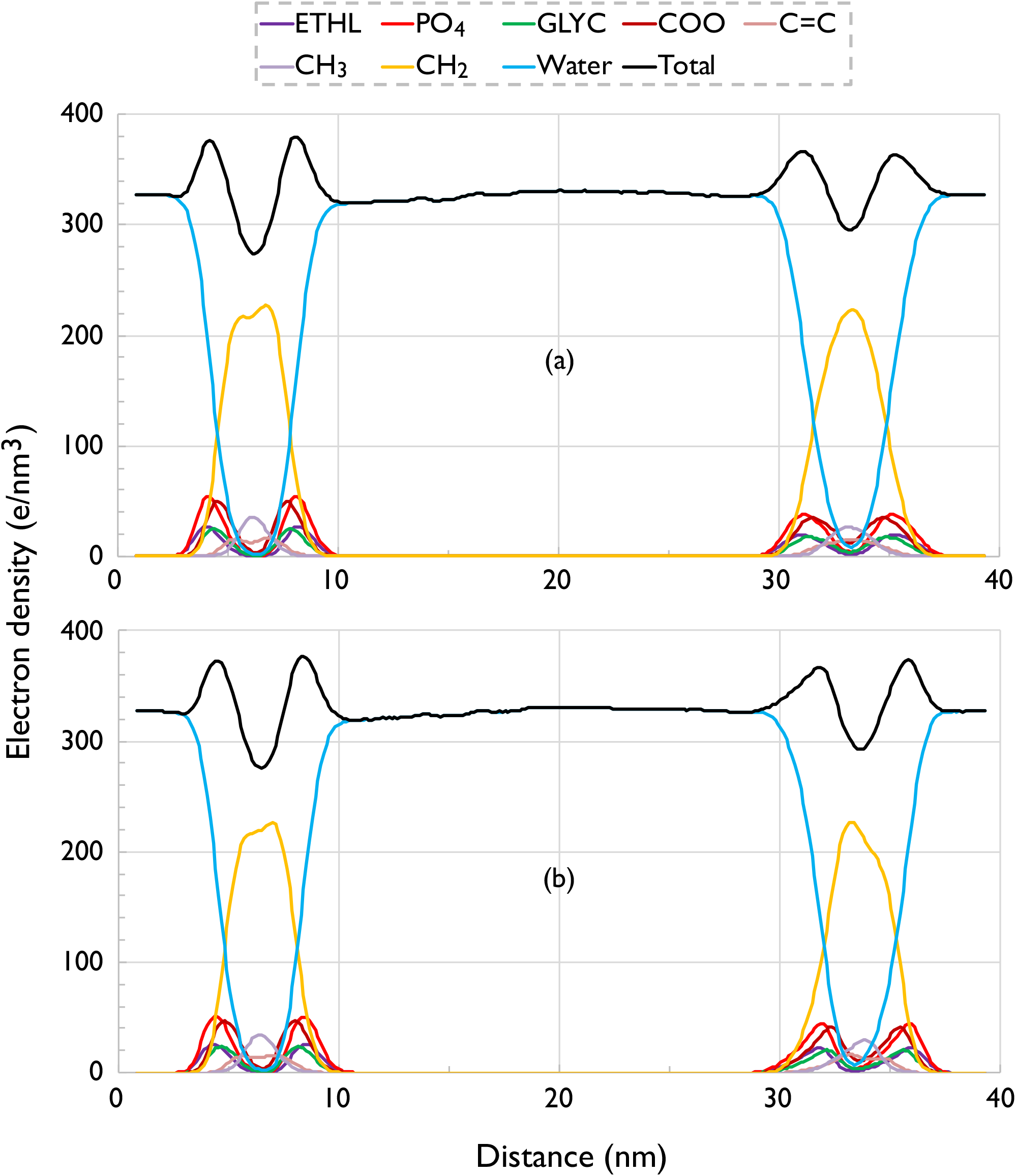
Electron density profiles along the membrane normals of the two trajectories from 50 to 400 ns from the MD simulations. The profiles were decomposed into the same groups as Figure 8.

Table 5 lists the bilayer thicknesses (D_HH_) of the inner and outer membranes obtained from the peak-to-peak distance of the total electron density in Figure 10. Trajectory (b) showed the same thickness between the inner and outer membranes, but trajectory (a) showed a thickness difference equal to 0.39 nm, which were time and space averages during 350 ns and in a square area with a side length of 40 nm. These were significantly large against the pure POPE bilayer simulations that showed only 0.1 nm thickness differences by the temperatures from 293 to 333 K. Nevertheless, it is remarkable that the membrane thickness for the small unit cell and large efflux system agreed, notwithstanding the large undulatory fluctuations of the membranes.

**Table 4:** Lipid bilayer thickness (D_HH_) derived from the peak-to-peak distance of the total electron density (Figure 8) and its standard errors in the tree trajectories.

| Lipid | DLPC | DMPC | DOPC | DPPC | POPC | SOPC | POPE |
| --- | --- | --- | --- | --- | --- | --- | --- |
| Simulation (nm) | 2.83 | 3.25 | 3.61 | 3.56 | 3.66 | 3.82 | 3.94 |
| Standard error (nm) | 0.05 | 0.08 | 0.06 | 0.05 | 0.02 | 0.03 | 0.06 |
| Experiment (nm) | 3.08<sup>41</sup> | 3.44<sup>61</sup> | 3.67<sup>44</sup> | 3.83<sup>39</sup> | 3.70<sup>42</sup> |  | 3.95<sup>40</sup> |
| Temperature (K) | 303 | 303 | 303 | 323 | 303 | 303 | 310 |

**Table 5:** Lipid bilayer thickness (D_HH_) derived from the peak-to-peak distance of the total electron density in Figure 10.

| Trajectory | (a) |  | (b) |  |
| --- | --- | --- | --- | --- |
| Membrane | inner | outer | inner | outer |
| Thickness (nm) | 3.86 | 4.25 | 3.97 | 3.97 |

The membrane thickness differences in trajectories (a) and (b) were caused by thermal fluctuations, which also produced the difference in the interaction energies between proteins and membranes (Figure 9). As previously mentioned, the Coul-SR difference between TolC-POPE and AcrB-POPE is larger in trajectory (a) than in trajectory (b). In trajectory (a), the Coul-SR of TolC-POPE became weaker, while the Coul-SR of AcrB-POPE became stronger. If the lipid volume is constant, the lipid area should decrease as the thickness increases. Since the outer membrane of (a) is thicker than the other membranes, its lipid area decreased so that the Coul-SR of TolC-POPE became weaker. Obviously, the membrane fluctuations affected the interactions with proteins. Therefore, the membrane protein dynamics should be studied by accurate lipid force fields. Currently, we are beginning to study the sophisticated efflux mechanism of the complex multidrug efflux pump.

## 5 Conclusions

The Lipid membrane is a typical inhomogeneous and anisotropic system that is not suitable for the traditional van der Waals cutoff scheme with the isotropic dispersion correction. Although the AMBER and CHARMM lipid force fields could produce the proper membrane characteristics using the traditional cutoff scheme without an additional surface tension, their lipid bilayers unacceptably shrank when using the LJ-PME method. We focused on the first-principles derivation of a new lipid force field. The point charges were obtained from the arithmetic averages of the RESP charges of many conformations of the head and tail parts. The torsion dihedral parameters of the 12 torsion angles of phospholipids were derived by a fast Fourier transform from the high-level *ab initio* torsion energy profiles calculated with df-LCCSD(T)/Aug-cc-pVTZ//df-LMP2/Aug-cc-pVTZ, which was previously used to obtain the torsion dihedral parameters of the protein backbone of the FUJI force field.^17^ These first-principles parameters were incorporated into a new lipid force field (FUJI) without any fittings to experimental data. Since the FUJI lipid force field is composed of modules—two heads (PC and PE) and five tails (LA, MY, OL, PA, and ST), it is easy to construct new lipid force fields from the modules.

The membrane characteristics of the new lipid force field were studied by MD simulations with the LJ-PME method and hydrogen virtual sites with small cells for seven pure lipid bilayers at different temperatures. The area per lipid and the lateral diffusion coefficients showed satisfactory agreement with experimental data, but the order parameters of the acyl tails showed significant differences from the experimental NMR order parameters and other simulations by traditional lipid force fields. Our results suggest some ambiguities regarding the scaling factor between the NMR order parameter and NMR observables such as the quadrupolar splitting ∆*ν_Q_* and effective dipolar coupling *d*_CH_. The electron density profiles along the membrane normal were calculated by the GROMACS density subprogram with the “-center” option for seven pure lipid bilayers, and the membrane thicknesses (D_HH_) agreed well with the experimental values.

We constructed the simulation system with the multidrug efflux transporter (AcrABZ-TolC) with inner and outer membranes using the new FUJI force field. Large undulatory fluctuations of membranes were always observed, but there were no disturbances in the membrane structure. The order parameters of the acyl tails were the same for the small and large membranes. Although the electron densities of the lipid parts were widely spread in comparison with those of the pure lipid bilayer, the average bilayer thickness (D_HH_) did not differ between the small and large membranes.

The new FUJI force field offers accurate modeling for complex systems consisting of various membrane proteins and lipids.

## 6 Acknowledgements

The author (H. F.) thanks Erik Lindahl for stimulating discussions that guided us in developing new lipid force fields to explore the anisotropic characteristics of membranes by the LJ-PME method. This work was supported partly by HPCI Strategic Program Field 1 (Project IDs: hp120297, hp130006, hp140228, and hp150230), the Innovative Drug Discovery Infrastructure through the Functional Control of Biomolecular Systems Priority Issue 1 in Post-K Supercomputer Development from MEXT (Project ID: hp160223, hp170255, and hp180191). We used the computational resources of the K computer through the HPCI system research project (Project ID: hp170318) and the computer in the Research Center for Computational Science, Okazaki. The author (K. S.) was supported partially by the Initiative on Promotion of Supercomputing for Young or Women Researchers, Information Technology Center in the University of Tokyo.

## References

(1) Dror, R. O.; Green, H. F.; Valant, C.; Borhani, D. W.; Valcourt, J. R.; Pan, A. C.; Arlow, D. H.; Canals, M.; Lane, J. R.; Rahmani, R.; Baell, J. B.; Sexton, P. M.; Christopoulos, A.; Shaw, D. E. Structural basis for modulation of a G-protein-coupled receptor by allosteric drugs. Nature 2013, 503, 295–299, DOI: 10.1038/nature12595.

(2) Ewald, P. P. Die Berechnung optischer und elektrostatischer Gitterpotentiale. Ann. Phys. 1921, 369, 253–287, DOI: 10.1002/andp.19213690304.

(3) Darden, T.; York, D.; Pedersen, L. Particle mesh Ewald: An *N* log(*N*) method for Ewald sums in large systems. J. Chem. Phys. 1993, 98, 10089–10092, DOI: 10.1063/1.464397.

(4) Allen, M. P.; Tildesley, D. J. Computer Simulation of Liquids; Oxford University Press: New York, 1987.

(5) Abraham, M.; Hess, B.; van der Spoel, D.; Lindahl, E. GROMACS Reference Manual. GROMACS Development teams: Royal Institute of Technology, Uppsala University, Sweden, 2016.

(6) Dickson, C. J.; Rosso, L.; Betz, R. M.; Walkercd, R. C.; Gould, I. R. GAFFlipid: a General Amber Force Field for the accurate molecular dynamics simulation of phospholipid. Soft Matter 2012, 8, 9617–9627, DOI: 10.1039/c2sm26007g.

(7) Jämbeck, J. P. M.; Lyubartsev, A. P. Derivation and Systematic Validation of a Refined All-Atom Force Field for Phosphatidylcholine Lipids. J. Phys. Chem. B 2012, 116, 3164–3179, DOI: 10.1021/jp212503e.

(8) Dickson, C. J.; Madej, B. D.; Skjevik, Å. A.; Betz, R. M.; Teigen, K.; Gould, I. R.; Walker, R. C. Lipid14: The Amber Lipid Force Field. J. Chem. Theory Comput. 2014, 10, 865–879, DOI: 10.1021/ct4010307.

(9) Klauda, J. B.; Venable, R. M.; Freites, J. A.; O’Connor, J. W.; Tobias, D. J.; Mondragon-Ramirez, C.; Vorobyov, I.; MacKerell Jr., A. D.; Pastor, R. W. Update of the CHARMM All-Atom Additive Force Field for Lipids: Validation on Six Lipid Types. J. Phys. Chem. B 2010, 114, 7830–7843, DOI: 10.1021/jp101759q.

(10) Wennberg, C. L.; Murtola, T.; Hess, B.; Lindahl, E. Lennard-Jones Lattice Summation in Bilayer Simulations Has Critical Effects on Surface Tension and Lipid Properties. J. Chem. Theory Comput. 2013, 9, 3527–3537, DOI: 10.1021/ct400140n.

(11) Wennberg, C. L.; Murtola, T.; Páll, S.; Abraham, M. J.; Hess, B.; Lindahl, E. Direct-Space Corrections Enable Fast and Accurate Lorentz-Berthelot Combination Rule Lennard-Jones Lattice Summation. J. Chem. Theory Comput. 2015, 11, 5737–5746, DOI: 10.1021/acs.jctc.5b00726.

(12) Berendsen, H. J. C.; van der Spoel, D.; van Drunen, R. GROMACS: A message-passing parallel molecular dynamics implementation. Comp. Phys. Comm. 1995, 91, 43–56, DOI: 10.1016/0010-4655(95)00042-E.

(13) Páll, S.; Abraham, M. J.; Kutzner, C.; Hess, B.; Lindahl, E. In Solving Software Challenges for Exascale; Markidis, S., Laure, E., Eds.; Lecture Notes in Computer Science; Springer International Publishing: Switzerland, 2015; Vol. 8750; pp 3–27, DOI: 10.1007/978-3-319-15976-8.

(14) Hess, B. P-LINCS: A Parallel Linear Constraint Solver for Molecular Simulation. J. Chem. Theory Comput. 2007, 4, 116–122, DOI: 10.1021/ct700200b.

(15) Fujitani, H.; Tanida, Y.; Matsuura, A. Massively parallel computation of absolute binding free energy with well-equilibrated states. Phys. Rev. E 2009, 79, 021914, DOI: 10.1103/PhysRevE.79.021914.

(16) Wang, J.; Wolf, R. M.; Caldwell, J. W.; Kollman, P. A.; Case, D. A. Development and testing of a general amber force field. J. Comput. Chem. 2004, 25, 1157–1174, DOI: 10.1002/jcc.20035.

(17) Fujitani, H.; Matsuura, A.; Sakai, S.; Sato, H.; Tanida, Y. High-Level ab Initio Calculations To Improve Protein Backbone Dihedral Parameters. J. Chem. Theory Comput. 2009, 5, 1155–1165, DOI: 10.1021/ct8005437.

(18) Grdadolnik, J.; Grdadolnik, S. G.; Avbelj, F. Determination of Conformational Preferences of Dipeptides Using Vibrational Spectroscopy. J. Phys. Chem. B 2008, 112, 2712–2718, DOI: 10.1021/jp7096313.

(19) Grdadolnik, J.; Mohacek-Grosev, V.; Baldwin, R. L.; Avbelj, F. Populations of the three major backbone conformations in 19 amino acid dipeptides. Proc. Natl. Acad. Sci. USA 2011, 108, 1794–1798, DOI: 10.1073/pnas.1017317108.

(20) Vymětal, J.; Vondrášek, J. The DF–LCCSD(T0) correction of the *ϕ/ψ* force field dihedral parameters significantly influences the free energy profile of the alanine dipeptide. Chem. Phys. Lett. 2011, 503, 301–304, DOI: 10.1016/j.cplett.2011.01.030.

(21) Tzanov, A. T.; Cuendet, M. A.; Tuckerman, M. E. How Accurately Do Current Force Fields Predict Experimental Peptide Conformations? An Adiabatic Free Energy Dynamics Study. J. Phys. Chem. B 2014, 118, 6539–6552, DOI: 10.1021/jp500193w.

(22) Frisch, M. J.; Trucks, G. W.; Schlegel, H. B.; Scuseria, G. E.; Robb, M. A.; Cheeseman, J. R.; Scalmani, G.; Barone, V.; Mennucci, B.; Petersson, G. A.; Nakatsuji, H.; Caricato, M.; Li, X.; Hratchian, H. P.; Izmaylov, A. F.; Bloino, J.; Zheng, G.; Sonnenberg, J. L.; Hada, M.; Ehara, M.; Toyota, K.; Fukuda, R.; Hasegawa, J.; Ishida, M.; Nakajima, T.; Honda, Y.; Kitao, O.; Nakai, H.; Vreven, T.; J. A. Montgomery, J.; Peralta, J. E.; Ogliaro, F.; Bearpark, M.; Heyd, J. J.; Brothers, E.; Kudin, K. N.; Staroverov, V. N.; Keith, T.; Kobayashi, R.; Normand, J.; Raghavachari, K.; Rendell, A.; Burant, J. C.; Iyengar, S. S.; Tomasi, J.; Cossi, M.; Rega, N.; Millam, J. M.; Klene, M.; Knox, J. E.; Cross, J. B.; Bakken, V.; Adamo, C.; Jaramillo, J.; Gomperts, R.; Stratmann, R. E.; Yazyev, O.; Austin, A. J.; Cammi, R.; Pomelli, C.; Ochterski, J. W.; Martin, R. L.; Morokuma, K.; Zakrzewski, V. G.; Voth, G. A.; Salvador, P.; Dannenberg, J. J.; Dapprich, S.; Daniels, A. D.; Farkas, O.; Foresman, J. B.; Ortiz, J. V.; Cioslowski, J.; Fox, D. J. Gaussian 09, Revision C.01. Gaussian, Inc.: Wallingford CT, 2010.

(23) Bayly, C. I.; Cieplak, P.; Cornell, W. D.; Kollman, P. A. A Well-Behaved Electrostatic Potential Based Method Using Charge Restraints for Deriving Atomic Charges: The RESP Model. J. Phys. Chem. 1993, 97, 10269–10280, DOI: 10.1021/j100142a004.

(24) Dupradeau, F.-Y.; Pigache, A.; Zaffran, T.; Savineau, C.; Lelong, R.; Grivel, N.; Lelong, D.; Rosanskia, W.; Cieplak, P. The R.E.D. tools: advances in RESP and ESP charge derivation and force field library building. Phys. Chem. Chem. Phys. 2010, 12, 7821–7839, DOI: 10.1039/c0cp00111b.

(25) Cornell, W. D.; Cieplak, P.; Bayly, C. I.; Gould, I. R.; Merz, Jr., K. M.; Ferguson, D. M.; Spellmeyer, D. C.; Fox, T.; Caldwell, J. W.; Kollman, P. A. A Second Generation Force Field for the Simulation of Proteins, Nucleic Acids, and Organic Molecules. J. Am. Chem. Soc. 1995, 117, 5179–5197, DOI: 10.1021/ja955032e.

(26) Wang, J.; Cieplak, P.; Kollman, P. A. How well does a restrained electrostatic potential (RESP) model perform in calculating conformational energies of organic and biological molecules? J. Comput. Chem. 2000, 21, 1049–1074, DOI: 10.1002/1096-987X(200009)21:12<1049::AID-JCC3>.

(27) Duan, Y.; Wu, C.; Chowdhury, S.; Lee, M. C.; Xiong, G.; Zhang, W.; Yang, R.; Cieplak, P.; Luo, R.; Lee, T.; Caldwell, J.; Wang, J.; Kollman, P. A Point-Charge Force Field for Molecular Mechanics Simulations of Proteins Based on Condensed-Phase Quantum Mechanical Calculations. J. Comput. Chem. 2003, 24, 1999–2012, DOI: 10.1002/jcc.10349.

(28) Hornak, V.; Abel, R.; Okur, A.; Strockbine, B.; Roitberg, A.; Simmerling, C. Comparison of Multiple Amber Force Fields and Development of Improved Protein Backbone Parameters. Proteins: Struct. Funct. Bioinf. 2006, 65, 712–725, DOI: 10.1002/prot.21123.

(29) Lindorff-Larsen, K.; Piana, S.; Palmo, K.; Maragakis, P.; Klepeis, J. L.; Dror, R. O.; Shaw, D. E. Improved side-chain torsion potentials for the Amber ff99SB protein force field. Proteins: Struct. Funct. Bioinf. 2010, 78, 1950–1958, DOI: 10.1002/prot.22711.

(30) Maier, J. A.; Martinez, C.; Kasavajhala, K.; Wickstrom, L.; Hauser, K. E.; Simmerling, C. ff14SB: Improving the Accuracy of Protein Side Chain and Backbone Parameters from ff99SB. J. Chem. Theory Comput. 2015, 11, 3696–713, DOI: 10.1021/acs.jctc.5b00255.

(31) Werner, H.-J.; Knowles, P. J.; Knizia, G.; Manby, F. R.; tz, M. S. Molpro: a general-purpose quantum chemistry program package. Comput. Mol. Sci. 2012, 2, 242–253, DOI: 10.1002/wcms.82.

(32) Werner, H. J.; Knowles, P. J.; Lindh, R.; Manby, F. R.; Schütz, M.; Celani, P.; Korona, T.; Mitrushenkov, A.; Rauhut, G.; Adler, T. B.; Amos, R. D.; Bernhardsson, A.; Berning, A.; Cooper, D. L.; Deegan, M. J. O.; Dobbyn, A. J.; Eckert, F.; Goll, E.; Hampel, C.; Hetzer, G.; Hrenar, T.; Knizia, G.; Köppl, C.; Liu, Y.; Lloyd, A. W.; Mata, R. A.; May, A. J.; McNicholas, S. J.; Meyer, W.; Mura, M. E.; Nicklass, A.; Palmieri, P.; Pflüger, K.; Pitzer, R.; Reiher, M.; Schumann, U.; Stoll, H.; Stone, A. J.; Tarroni, R.; Thorsteinsson, T.; Wang, M.; Wolf, A. MOLPRO. Cardiff School of Chemistry, Cardiff University: Cardiff, UK, 2006.

(33) Wu, E. L.; Cheng, X.; Jo, S.; Rui, H.; Song, K. C.; avila Contreras, E. M. D.; Qi, Y.; Lee, J.; Monje-Galvan, V.; Venable, R. M.; Klauda, J. B.; Im, W. CHARMM-GUI Membrane Builder Toward Realistic Biological Membrane Simulations. J. Comput. Chem. 2014, 35, 1997–2004, DOI: 10.1002/jcc.23702.

(34) Nosé, S. A molecular dynamics method for simulations in the canonical ensemble. Mol. Phys. 1984, 52, 255–268, DOI: 10.1080/00268978400101201.

(35) Hoover, W. G. Canonical dynamics: Equilibrium phase-space distributions. Phys. Rev. A 1985, 31, 1695–1697, DOI: 0.1103/PhysRevA.31.1695.

(36) Berendsen, H. J. C.; Postma, J. P.; DiNola, A.; Haak, J. R. Molecular dynamics with coupling to an external bath. J. Chem. Phys. 1984, 81, 3684–3690, DOI: 10.1063/1.448118.

(37) Parrinello, M.; Rahman, A. Polymorphic transitions in single crystals: A new molecular dynamics method. J. Appl. Phys. 1981, 52, 7182–7190, DOI: 10.1063/1.328693.

(38) Petrache, H. I.; Dodd, S. W.; Brown, M. F. Area per Lipid and Acyl Length Distributions in Fluid Phosphatidylcholines Determined by 2H NMR Spectroscopy. Biophys. J 2000, 79, 3172–2192, DOI: 10.1016/S0006-3495(00)76551-9.

(39) Nagle, J. F.; Tristram-Nagle, S. Structure of lipid bilayers. Biochim. Biophs. Acta 2000, 1469, 159–195, DOI: 10.1016/S0304-4157(00)00016-2.

(40) Rappolt, M.; Hickel, A.; Bringezu, F.; Lohner, K. Mechanism of the Lamellar/Inverse Hexagonal Phase Transition Examined by High Resolution X-Ray Diffraction. Biophys. J 2003, 84, 3111–3122, DOI: 10.1016/S0006-3495(03)70036-8.

(41) Kučerka, N.; Liu, Y.; Chu, N.; Petrache, H. I.; Tristram-Nagle, S.; Nagle, J. F. Structure of Fully Hydrated Fluid Phase DMPC and DLPC Lipid Bilayers Using X-Ray Scattering from Oriented Multilamellar Arrays and from Unilamellar Vesicles. Biophys. J 2005, 88, 2626–2637, DOI: 10.1529/biophysj.104.056606.

(42) Kučerka, N.; Tristram-Nagle, S.; Nagle, J. F. Structure of Fully Hydrated Fluid Phase Lipid Bilayers with Monounsaturated Chains. J. Membrane Biol. 2006, 208, 193–202, DOI: 10.1007/s00232-005-7006-8.

(43) Kučerka, N.; Nagle, J. F.; Sachs, J. N.; Feller, S. E.; Pencer, J.; Jackson, A.; Katsaras, J. Lipid Bilayer Structure Determined by the Simultaneous Analysis of Neutron and X-Ray Scattering Data. Biophys. J 2008, 95, 2356–2367, DOI: 10.1529/biophysj.108.132662.

(44) Pan, J.; Tristram-Nagle, S.; Kučerka, N.; Nagle, J. F. Temperature Dependence of Structure, Bending Rigidity, and Bilayer Interactions of Dioleoylphosphatidylcholine Bilayers. Biophys. J 2008, 94, 117–124, DOI: 10.1529/biophysj.107.115691.

(45) Kučerka, N.; Nieh, M.-P.; Katsaras, J. Fluid phase lipid areas and bilayer thicknesses of commonly used phosphatidylcholines as a function of temperature. BBA-Biomembranes 2011, 1808, 2761–2771, DOI: 10.1016/j.bbamem.2011.07.022.

(46) Kučerka, N.; van Oosten, B.; JianjunPan,; Heberle, F. A.; Harroun, T. A.; Katsaras, J. Molecular Structures of Fluid Phosphatidylethanolamine Bilayers Obtained from Simulation-to-Experiment Comparisons and Experimental Scattering Density Profiles. J. Phys. Chem. B 2015, 119, 1947–1956, DOI: 10.1021/jp511159q.

(47) Vaz, W. L. C.; Clegg, R. M.; Hallmann, D. Translational diffusion of lipids in liquid crystalline phase phosphatidylcholine multibilayers. A comparison of experiment with theory. Biochemistry 1985, 24, 781–786, DOI: 10.1021/bi00324a037.

(48) Almeida, P. F. F.; Vaz, W. L. C.; Thompson, T. E. Lateral Diffusion in the Liquid Phases of Dimyristoylphosphatidylcholine/Cholesterol Lipid Bilayers: A Free Volume Analysis. Biochemistry 1992, 31, 6739–6747, DOI: 10.1021/bi00144a013.

(49) Ratto, T. V.; Longo, M. L. Obstructed Diffusion in Phase-Separated Supported Lipid Bilayers: A Combined Atomic Force Microscopy and Fluorescence Recovery after Photobleaching Approach. Biophys. J 2002, 83, 3380–3392, DOI: 10.1016/S0006-3495(02)75338-1.

(50) Kuśba, J.; Li, L.; Gryczynski, I.; Piszczek, G.; Johnson, M.; Lakowicz, J. R. Lateral Diffusion Coefficients in Membranes Measured by Resonance Energy Transfer and a New Algorithm for Diffusion in Two Dimensions. Biophys. J 2002, 82, 1358–1372, DOI: 10.1016/S0006-3495(02)75491-X.

(51) Filippov, A.; Orädd, G.; Lindblom, G. Influence of Cholesterol and Water Content on Phospholipid Lateral Diffusion in Bilayers. Langmuir 2003, 19, 6397–6400, DOI: 10.1021/la034222x.

(52) Scheidt, H. A.; Huster, D.; Gawrisch, K. Diffusion of Cholesterol and Its Precursors in Lipid Membranes Studied by 1H Pulsed Field Gradient Magic Angle Spinning NMR. Biophys. J 2005, 89, 2504–2512, DOI: 10.1529/biophysj.105.062018.

(53) Filippov, A.; Orädd, G.; Lindblom, G. Domain Formation in Model Membranes Studied by Pulsed-Field Gradient-NMR: The Role of Lipid Polyunsaturation. Biophys. J 2007, 93, 3182–3190, DOI: 10.1529/biophysj.107.111534.

(54) Warschawski, D. E.; Devaux, P. F. Order parameters of unsaturated phospholipids in membranes and the effect of cholesterol: a 1H-13C solid-state NMR study at natural abundance Order parameters of unsaturated phospholipids in membranes and the effect of cholesterol: a 1H–13C solid-state NMR study at natural abundance. Eur. Biophys. J. 2005, 34, 987–996, DOI: 10.1007/s00249-005-0482-z.

(55) Ferreira, T. M.; Coreta-Gomes, F.; Ollila, O. H. S.; Moreno, M. J.; Vaz, W. L. C.; Topgaard, D. Cholesterol and POPC segmental order parameters in lipid membranes: solid state 1H–13C NMR and MD simulation studies. Phys. Chem. Chem. Phys. 2013, 15, 1976–1989, DOI: 10.1039/c2cp42738a.

(56) Shaikh, S. R.; Brzustowicz, M. R.; Gustafson, N.; Stillwell, W.; Wassall, S. R. Monounsaturated PE Does Not Phase-Separate from the Lipid Raft Molecules Sphingomyelin and Cholesterol: Role for Polyunsaturation? Biochemistry 2002, 41, 10593–10602, DOI: 10.1021/bi025712b.

(57) Berger, O.; Edholm, O.; Jahnig, F. Molecular Dynamics Simulations of a Fluid Bilayer of Dipalmitoylphosphatidylcholine at Full Hydration, Constant Pressure, and Constant Temperature. Biophys. J 1997, 72, 2002–2013, DOI: 10.1016/S0006-3495(97)78845-3.

(58) Piggot, T. J.; Allison, J. R.; Sessions, R. B.; Essex, J. W. On the Calculation of Acyl Chain Order Parameters from Lipid Simulations. J. Chem. Theory Comput. 2017, 13, 5683–5696, DOI: 10.1021/acs.jctc.7b00643.

(59) Ollila, O. H. S.; Miettinen, M. S.; Vogel, A. Accuracy of order parameter measurements. The NMRlipids Collaboration, 2015.

(60) Rosso, L.; Gould, I. R. Structure and Dynamics of Phospholipid Bilayers Using Recently Developed General All-Atom Force Fields. J. Comput. Chem. 2008, 29, 24–27, DOI: 10.1002/jcc.20675.

(61) Petrache, H. I.; Tristram-Nagle, S.; Nagle, J. F. Fluid phase structure of EPC and DMPC bilayers. Chem. Phys. Lipids 1998, 95, 83–94, DOI: 10.1016/S0009-3084(98)00068-1.

(62) Wang, Z.; Fan, G.; Hryc, C. F.; Blaza, J. N.; Serysheva, I. I.; Schmid, M. F.; Chiu, W.; Luisi, B. F.; Du, D. An allosteric transport mechanism for the AcrAB-TolC multidrug efflux pump. eLife 2017, 6, e24905, DOI: 10.7554/eLife.24905.

(63) Schmidt, T. H.; Kandt, C. LAMBADA and InflateGRO2: Efficient Membrane Alignment and Insertion of Membrane Proteins for Molecular Dynamics Simulations. J. Chem. Inf. Model. 2012, 52, 2657–2669, DOI: 10.1021/ci3000453.

(64) Beglov, D.; Roux, B. Finite representation of an infinite bulk system: Solvent boundary potential for computer simulations. J. Chem. Phys. 1994, 100, 9050–9063, DOI: 10.1063/1.466711.

(65) Roux, B. Valence Selectivity of the Gramicidin Channel: A MolecularDynamics Free Energy Perturbation Study. Biophys. J 1996, 71, 3177–3185, DOI: 10.1016/S0006-3495(96)79511-5.

